# Transcriptional regulation by NR5A2 couples cell differentiation and inflammation in the pancreas

**DOI:** 10.1101/209411

**Authors:** Isidoro Cobo, Paola Martinelli, Marta Flández, Latifa Bakiri, Mingfeng Zhang, Enrique Carrillo-de-Santa-Pau, Jinping Jia, Liv Thommesen, Torunn Bruland, Natalia del Pozo, Sara Olson, Jill Smith, William R. Bamlet, Gloria M Petersen, Núria Malats, Laufey T Amundadottir, Erwin F. Wagner, Francisco X Real

## Abstract

Tissue-specific differentiation and inflammatory programmes are thought to independently contribute to disease. The orphan nuclear receptor NR5A2 is a key regulator of pancreas differentiation and SNPs in or near the human gene are associated with risk of pancreatic cancer. In mice, *Nr5a2* heterozygosity sensitizes the pancreas to damage, impairs regeneration, and cooperates with mutant KRas in tumor progression. Through global transcriptomic analysis, we uncover a basal pre-inflammatory state in the pancreas of *Nr5a2* heterozygous mice that is reminiscent of pancreatitis-induced inflammation and is conserved in histologically normal human pancreata with reduced NR5A2 mRNA expression. In *Nr5a2*^+/−^ mice, Nr5a2 undergoes a dramatic transcriptional switch relocating from tissue-specific to inflammatory loci thereby promoting AP-1-dependent gene transcription. Importantly, deletion of c-*Jun* in the pancreas of these mice rescues the pre-inflammatory phenotype and the defective regenerative response to damage. These findings provide compelling evidence that the same transcriptional networks supporting homeostasis in normal tissue can be subverted to foster inflammation upon genetic or environmental constraints.

## Introduction

Inflammatory responses contribute to the restoration of tissue integrity after insults that disrupt homeostasis ^1^. However, when persistent, they compromise tissue-resident cell function and are a risk factor for cancer in a variety of epithelial tissues, including the pancreas. The current notion is that the control of inflammatory responses relies on transcriptional networks distinct from those involved in cell differentiation ^2,3^.

The orphan nuclear receptor NR5A2 (also known as LRH-1) participates in a wide variety of processes including cholesterol and glucose metabolism in the liver, resolution of ER stress, intestinal glucocorticoid production, pancreatic development, and acinar differentiation ^4–8^. In mice, *Nr5a2* deletion leads to variable degrees of pancreatic hypoplasia depending on the timing of its inactivation ^6,8^. By contrast, constitutive *Nr5a2* heterozygosity allows normal pancreatic development but is associated with impaired regeneration upon induction of mild acute inflammation and sensitizes pancreatic epithelial cells to mutant KRas-driven preneoplastic lesions and pancreatic ductal adenocarcinoma (PDAC) ^9^.

In humans, single nucleotide polymorphisms (SNPs) in the vicinity of *NR5A2* have been associated with PDAC risk through genome wide association studies (GWAS) ^10,11^. While the mechanism(s) underlying risk have not been elucidated, NR5A2 expression has recently been proposed to define a subtype of PDAC with mixed exocrine and endocrine features ^12^.

Here, we show that *Nr5a2* heterozygosity leads to an epithelial cell-autonomous pre-inflammatory state in the pancreas that is largely recapitulated in histologically normal human pancreas in association with low NR5A2 transcript levels. In the pancreas of *Nr5a2*^+/−^ mice, and in wild type mice upon inflammatory stimuli, Nr5a2 undergoes a transcriptional switch and relocates from promoters of pancreatic differentiation genes to those of inflammatory genes through genetic and biochemical interaction with components of the AP-1 family. The phenotype of *Nr5a2*^+/−^ mice is rescued by deletion of *c-Jun* in the pancreas. We provide compelling evidence that the same transcriptional networks that support homeostasis in normal tissue can be subverted to foster inflammation upon genetic or environmental constraints.

## Results

### NR5A2 expression, PDAC risk variants at chr1q32.1, and pancreatitis

The cooperation between germline *Nr5a2* haploinsufficiency, a somatic oncogenic mutation, and pancreatic inflammation to promote pancreatic cancer suggested the existence of functional interactions relevant to human PDAC ^9^. We therefore analyzed the relationship between NR5A2 expression and pancreatitis in patients with PDAC. In two independent case series, the group of tumors with a lower NR5A2 expression was significantly enriched in subjects with self-reported personal history of chronic pancreatitis (combined analysis of both patient series, *P*=0.001) (Extended Data Table 1). We assessed the association between SNPs associated with PDAC risk ^10^ and NR5A2 protein levels using immunohistochemistry (IHC) on 110 PDAC tissue samples and germline genotyping from the same subjects. We noted suggestive associations between the risk-increasing allele of rs3790844 (T) and lower NR5A2 protein levels, both using average histoscores (*P*=0.097, β=−18.0) and average histoscore quantiles (*P*=0.028, β=−0.57) (Extended Data Figure 1). While these findings do not establish these SNPs as functional variants at this risk locus, they suggest that lower levels of NR5A2 may underlie susceptibility to PDAC in carriers of risk alleles at chr1q32.1.

### Low NR5A2 expression in normal pancreas is associated with a pre-inflammatory phenotype in mice and humans

How *Nr5a2* heterozygosity predisposes to impaired pancreatic regeneration and cancer is not known. The pancreas of adult *Nr5a2*^+/−^ mice is histologically normal, as is the expression of mRNAs coding for transcription factors involved in acinar differentiation (*Ptf1a, Rbpjl*) and digestive enzymes (*Cpa*, *Cel*, *Ela1b*, *Ctrb1*, and *Pnlip*) (Extended Data Figure 2A). Moreover, expression of Ptf1a, a master regulator of pancreatic development and differentiation, and its target Cpa was similar in *Nr5a2* wild type and heterozygous mice (Extended Data Figure 2A,B). We hypothesized that a subtle defect might exist in basal conditions because *Nr5a2*^+/−^ mice recovered poorly from a mild acute pancreatitis ^9^ and used RNA-Seq to compare the transcriptome of pancreata from 8-10 week old *Nr5a2*^+/+^ and *Nr5a2*^+/−^ mice.

A total of 929 genes and 101 genes were significantly up-regulated or down-regulated, respectively, in basal conditions in *Nr5a2*^+/−^ pancreata. Gene Set Enrichment Analysis (GSEA) revealed that 21 of the 23 gene sets identified as over-represented in *Nr5a2*^+/−^ mice belong to inflammation-related pathways (Extended Data Table 2 and Figure 1A). Sixty-eight percent of the up-regulated genes belong to inflammatory pathways (Extended Data Table 3). Among them are those coding for cytokines (*Ccl5, Ccl7, Ccl9, Cxcl10, Cxcl16*), complement components (*C1qb, C3, C5, Cfb, Cfd, Cfp, C3ar1, C5ar1*), and metalloproteases (*Mmp7, Mmp9*). RNA-Seq results were confirmed by quantitative RT-PCR (RT-qPCR) (Figure 1B), thus suggesting a basal pre-inflammatory status in *Nr5a2*^+/−^ pancreas. In agreement with the expression profiles, the promoters of genes that were up-regulated in *Nr5a2*^+/−^ pancreata showed increased histone marks associated with active chromatin, such as H3K27ac, and decreased marks associated with repressive chromatin, such as H3K27me3 (Extended Data Figure 3). The inverse results were obtained when the promoters of down-regulated genes were analysed (data not shown).

**Figure 1.**
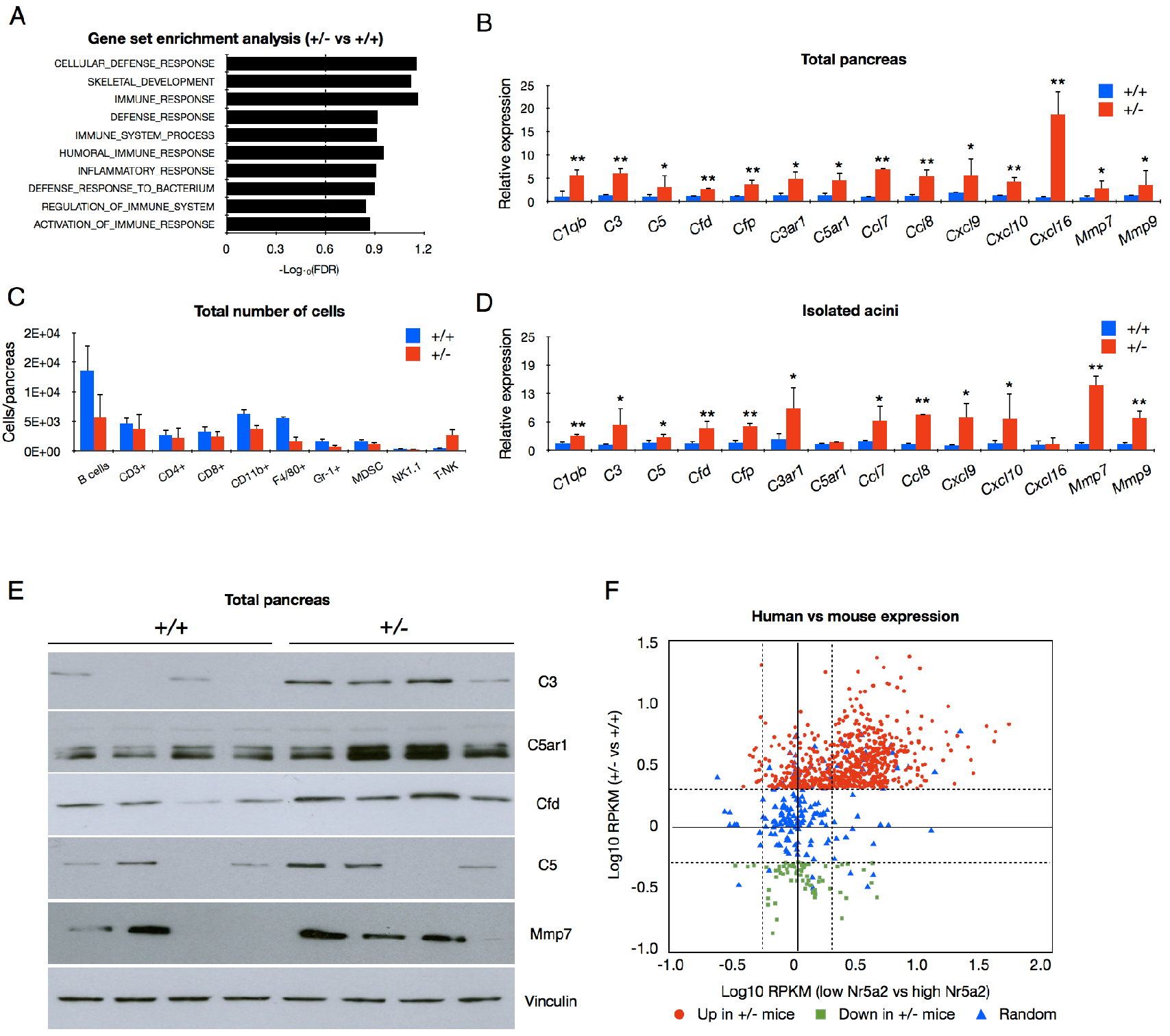
Reduced Nr5a2 expression is associated with a pre-inflammatory state in the normal pancreas of mice and humans. (A) Gene set enrichment analysis performed on the genes differentially expressed in *Nr5a2*^+/−^ (n=3) versus wild type mice (n=3). (B) RT-qPCR showing up-regulation of inflammatory genes in *Nr5a2*^+/−^ pancreata (relative to wild type) (n=5 for each condition). (C) FACS analysis of the indicated populations of inflammatory cells in the pancreas of wild type and *Nr5a2*^+/−^ mice (n=4/group). (D) RT-qPCR showing the up-regulation of inflammatory genes in primary acinar cells from *Nr5a2*^+/−^ (relative to wild type) (n=4/group). (E) Western blots showing up-regulation of inflammatory proteins in the pancreas of *Nr5a2*^+/−^ mice (n=4/group). (F) Scatter plot showing the relationship between the expression of up-regulated, down-regulated, or a random set of genes in control *Nr5a2*^+/+^ vs. *Nr5a2*^+/−^ mice (y-axis) and in histologically normal human pancreatic tissue samples (x-axis, top high vs. low quartiles of NR5A2 mRNA expression, as determined by RNA-Seq analysis).

Acinar cells constitute approximately 90% of epithelial cells in normal pancreas, while inflammatory cells are very scarce. We did not find significant differences in the total number of Cd45^+^ cells, nor in the proportion of specific immune cell subpopulations, in the pancreas of wild type and *Nr5a2*^+/−^ mice (Figure 1C; Extended Data Figure 2C), ruling out that the activation of an inflammatory programme resulted from increased leukocyte infiltration. In addition, we isolated acinar cells and confirmed by RT-qPCR the up-regulation of inflammatory transcripts (Figure 1D). Western blotting analyses also showed an increased expression of C3, C5ar1, Cfd, C5, and Mmp7 in extracts from both total pancreas and isolated acinar cells (Figure 1E and Extended Data Figure 4A). Moreover, IHC analyses confirmed that C5ar1 and Cfd are expressed in acinar cells (Extended Data Figure 2D). These results indicate that *Nr5a2* haploinsufficiency leads to a pre-inflammatory gene expression profile in pancreatic epithelial cells.

To determine whether the genes identified as being differentially expressed in the pancreas of *Nr5a2*^+/−^ mice were similarly deregulated in human pancreas, we performed RNA-Seq of 95 histologically normal human pancreatic tissue samples (Zhang et al, unpublished data). We then compared transcript levels in samples with *NR5A2* expression in the top quartile (high, n=24) with those in the bottom quartile (low, n=24). We assessed three gene sets from the mouse transcriptome analysis: genes up-regulated in *Nr5a2*^+/−^ pancreata (n=725), genes down-regulated in *Nr5a2*^+/−^ pancreata (n=67), and a random set of genes (n=249). Seventy-eight percent of genes whose expression was up-regulated in *Nr5a2*^+/−^ mice were differentially expressed in human pancreatic samples with low vs. high *NR5A2* expression with a nominal *P* value <0.05). Of these, 92% were significantly up-regulated. On the other hand, 44% of the genes down-regulated in *Nr5a2*^+/−^ mice were differentially represented in samples with low vs. high *NR5A2* expression with a P<0.05 and only 50% of them were also down-regulated. Among the group of randomly selected genes, 44% were differentially expressed in samples with low vs. high *NR5A2* expression with a P<0.05 (Figure 1F). Overall, the genes that were up-regulated in the mouse were also up-regulated in the NR5A2^low^ vs. the NR5A2^high^ human pancreas samples, when compared with the random list of genes (P=10^−31^). By contrast, the down-regulated genes did not follow a concordant pattern in mouse and human pancreas (P=0.58). These results support the notion that NR5A2-dependent expression of inflammatory genes is conserved between murine and human pancreatic tissues and provide a framework to decipher the relationship between *NR5A2* genotype, expression, and PDAC risk using mouse models.

### *Nr5a2* heterozygosity is associated with increased AP-1 binding to inflammatory genes

To assess whether Nr5a2 directly regulates the pre-inflammatory state, we took advantage of published pancreatic Nr5a2 ChIP-seq datasets (SRR389293 and SRR389294) ^6^. Interestingly, the promoters of 89% of the genes whose expression was up-regulated in *Nr5a2*^+/−^ mice contain putative Nr5a2 binding sites but only 7% exhibited binding of Nr5a2 within 2.5 Kb from the transcriptional start site (TSS), suggesting the participation of an indirect mechanism (Figure 2A). Promoter Scanning Analysis (PScan) of up-regulated genes showed significant enrichment of binding motifs for several transcription factors, including AP-1 and NF-kB. Similar enrichment of AP-1 and NF-kB motifs was found when the list of up-regulated genes was computed with GSEA using the MSigDB C3 transcription factor target gene set collection (not shown).

**Figure 2.**
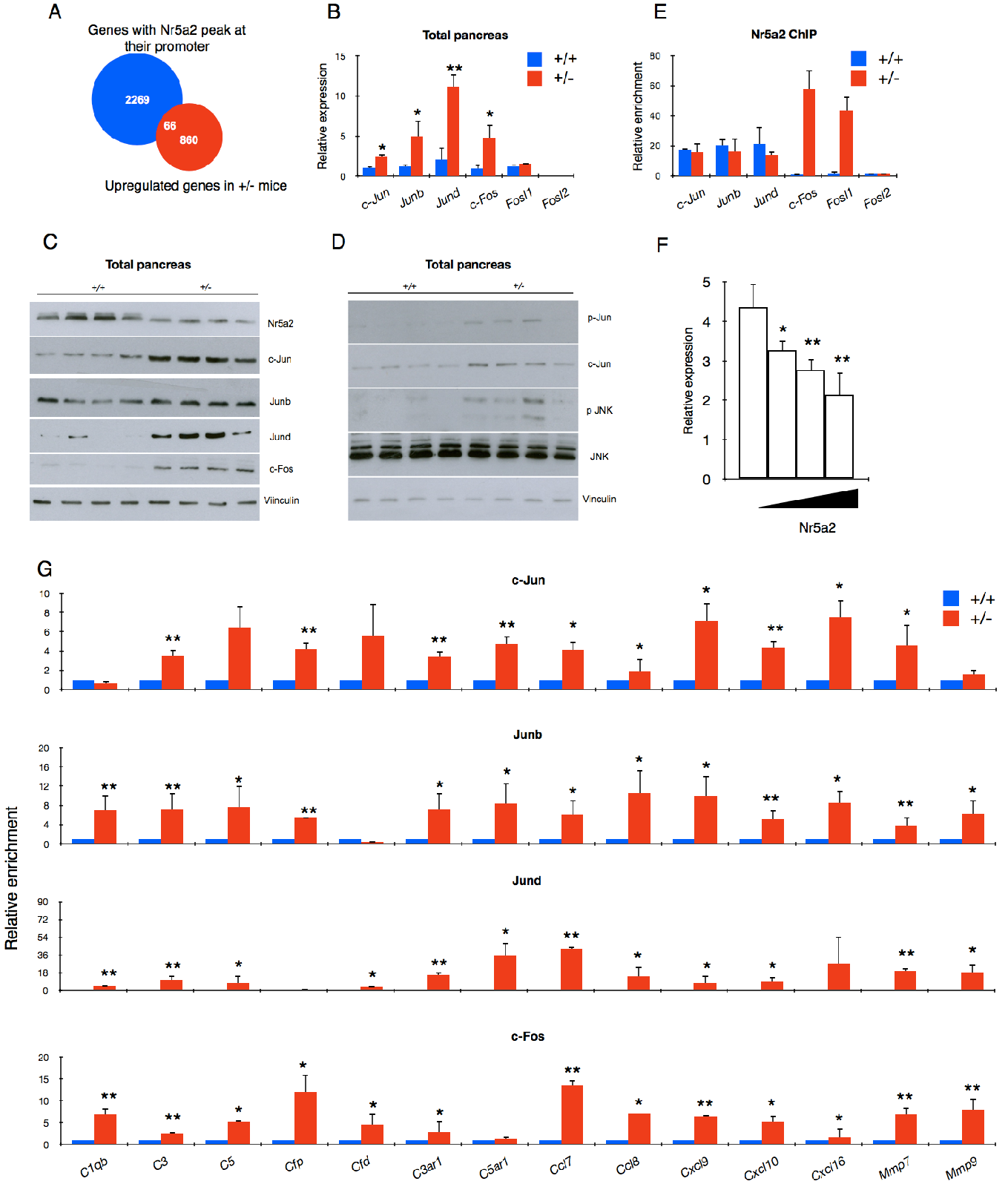
AP-1 components are up-regulated, and bind to the promoter of inflammatory genes, in the pancreas of *Nr5a2*^+/−^ mice. (A) A minor fraction of the promoters of genes up-regulated in the pancreas of *Nr5a2*^+/−^ mice display Nr5a2 peaks in ChIP-Seq experiments using normal pancreas. (B) RT-qPCR analysis of AP-1 component expression in wild type and *Nr5a2*^+/−^ mice (n=6/group). (C) Western blots showing expression of NR5A2 and AP-1 components in total pancreas of wild type and *Nr5a2*^+/−^ mice (n=4/group). (D) Western blots showing expression of p-Jun and p-Jnk in total pancreas of wild type and *Nr5a2*^+/−^ mice (n=4/group). (E) Nr5a2 enrichment at the promoter of AP-1 genes using ChIP-qPCR (n=7/group). (F) Nr5a2 overexpression in 293 cells leads to reduced c-Jun mRNA expression. Data correspond to the mean of 5 independent experiments. (G) ChIP-qPCR analysis of the occupancy of the promoter of inflammatory genes by AP-1 components in wild type and *Nr5a2*^+/−^ mice (n=5/group).

We next investigated the potential implication of the AP-1 family members in the basal inflammatory phenotype found in *Nr5a2*^+/−^ pancreata. c-Jun, Junb, Jund, and c-Fos were found significantly up-regulated in *Nr5a2*^+/−^ pancreata at the mRNA (Figure 2B) and protein (Figure 2C) levels. We confirmed that these changes occurred in epithelial cells using freshly isolated acini (Extended Data Figure 4B) and IHC (results for c-Jun shown in Extended Data Figure 5). Importantly, increased p-Jun and p-Jnk expression was observed in *Nr5a2*^+/−^ pancreata (Figure 2D). Using chromatin immunoprecipitation (ChIP-PCR), we found that Nr5a2 binds similarly the promoter of *c-Jun, Junb,* and *Jund* in the pancreas of wild type and *Nr5a2*^+/−^ mice. By contrast, Nr5a2 showed significantly increased binding to the promoter of *c-Fos* and Fos-like1 (*Fosl1*) genes in *Nr5a2*^+/−^ pancreata (Figure 2E). Ectopic Nr5a2 expression in HEK 293 cells led to a dose-dependent selective decrease of c-Jun mRNA levels (Figure 2F), indicating that changes in Nr5a2 levels can lead to the modulation of c-Jun expression.

To determine whether AP-1 could mediate the up-regulation of inflammation-related genes, we performed ChIP-qPCR for *c-Jun*, *c-Fos*, *Junb*, and *Jund* in mouse pancreatic tissue and showed that *Nr5a2*^+/−^ mice display an increased binding of these proteins to the promoters of up-regulated inflammatory genes (Figure 2G). Importantly, we did not observe increased recruitment of AP-1 to the promoter of down-regulated genes (not shown).

### *Nr5a2* haploinsufficiency mimics acute pancreatitis

The basal pre-inflammatory gene expression profile of *Nr5a2*^+/−^ mice suggests a subclinical pancreatitis-like state. To explore this notion, we analyzed the pancreatic transcriptome of wild type and *Nr5a2*^+/−^ mice 8, 24, and 48 h after induction of a mild acute pancreatitis (7 hourly doses of caerulein) using RNA-Seq. Principal Component Analysis (PCA) highlighted the divergence of the transcriptomes of *Nr5a2*^+/−^ and wild type pancreata in basal conditions (0 h). These differences were completely, but transiently, eroded at early stages of pancreatitis (8 h) (Figure 3A). The transcriptome of *Nr5a2*^+/−^ pancreata in basal conditions was highly similar to that of wild type mice 8h after the induction of pancreatitis (Figure 3A). When the set of genes up-regulated in *Nr5a2*^+/−^ mice in basal conditions was considered, the differences between the *Nr5a2*^+/−^ and the *Nr5a2*^+/+^ pancreata were notably reduced 8-24 h after induction of pancreatitis (Figure 3B). A similar, but inverse, dynamic pattern was observed with the set of down-regulated genes (Figure 3C). Expression of a random set of genes was unaffected (Figure 3D). These results indicate that, globally, the basal transcriptome of the pancreas of *Nr5a2*^+/−^ mice resembles that of an early acute pancreatitis state.

**Figure 3.**
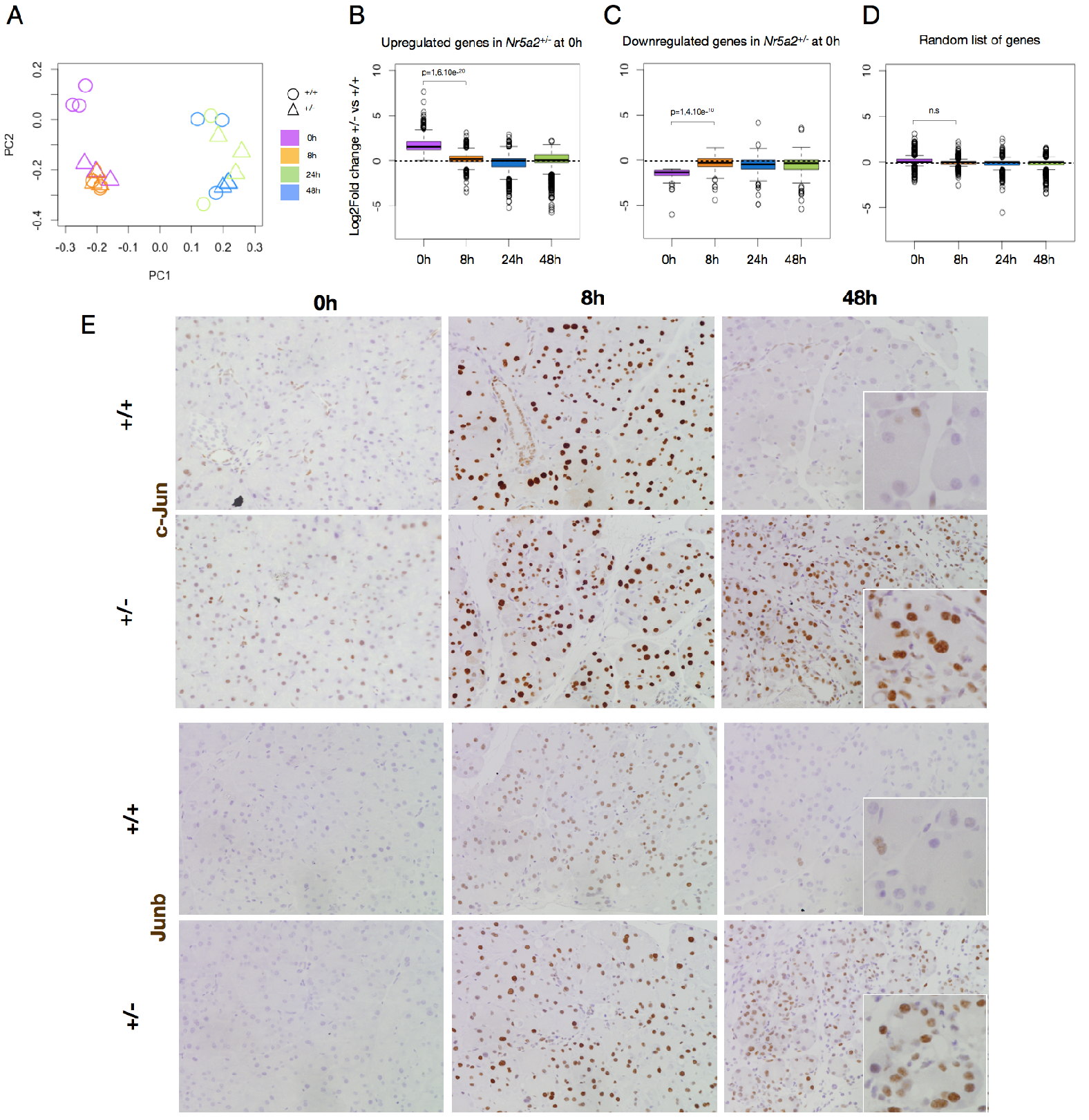
*Nr5a2* haploinsufficiency causes a basal pre-inflammatory state similar to that associated with early stages of pancreatitis. (A) Principal component analysis of the transcriptome of wild type and *Nr5a2*^+/−^ mice in basal conditions and 8, 24, and 48 h after the initiation of a caerulein-induced pancreatitis (7 hourly injections). (B-D) Comparative expression (wild type vs. *Nr5a2*^+/−^ mice) of the up-regulated (B), down-regulated (C), or control (D) genes over time after induction of pancreatitis. (E) Immunohistochemical analysis shows persistent over-expression of AP-1 components during the recovery period after induction of acute pancreatitis (see also Extended Data Figure 6) (n=5/group; representative results are shown).

Consistent with the results described above, we observed a strong, and similar, up-regulation of expression of AP-1 proteins at early time points during pancreatitis (8h) in mice of both genotypes. However, at later time points (24 h and 48 h), persistent up-regulation of AP-1 proteins was observed only in *Nr5a2*^+/−^ mice (Figure 3E and Extended Data Figure 6).

The repeated dosing of caerulein used to induce a standard acute pancreatitis hampers the interpretation of the dynamics of acute signalling/transcriptional responses. Therefore, we analyzed the effects of the administration of a single dose of caerulein that does not induce histological changes or inflammatory cell infiltration (Extended Data Figure 7A). Immunohistochemical analyses showed that c-Jun, c-Fos, Jund, Fra1, and Fra2 were up-regulated to a similar extent in both *Nr5a2*^+/+^ and *Nr5a2*^+/−^ pancreata 30 min after caerulein administration (Extended Data Figure 7 and data not shown). Up-regulation of c-Jun, c-Fos, Jund, Fra1 and Fra2 and c-Jun phosphorylation preceded the phosphorylation of Stat3 observed at 2 h (Extended Data Figure 7B).

We analyzed by RT-qPCR the expression of a subset of the genes differentially overexpressed in *Nr5a2*^+/−^ pancreata in basal conditions. A single dose of caerulein failed to cause histological pancreatitis but was sufficient to promote by 30-60 min, in the pancreas of wild type mice, an inflammatory profile that highly resembles that of *Nr5a2*^+/−^ pancreata. A complete resolution of this expression profile could be demonstrated in wild type mice by 12 h (Figure 4A, B).

**Figure 4.**
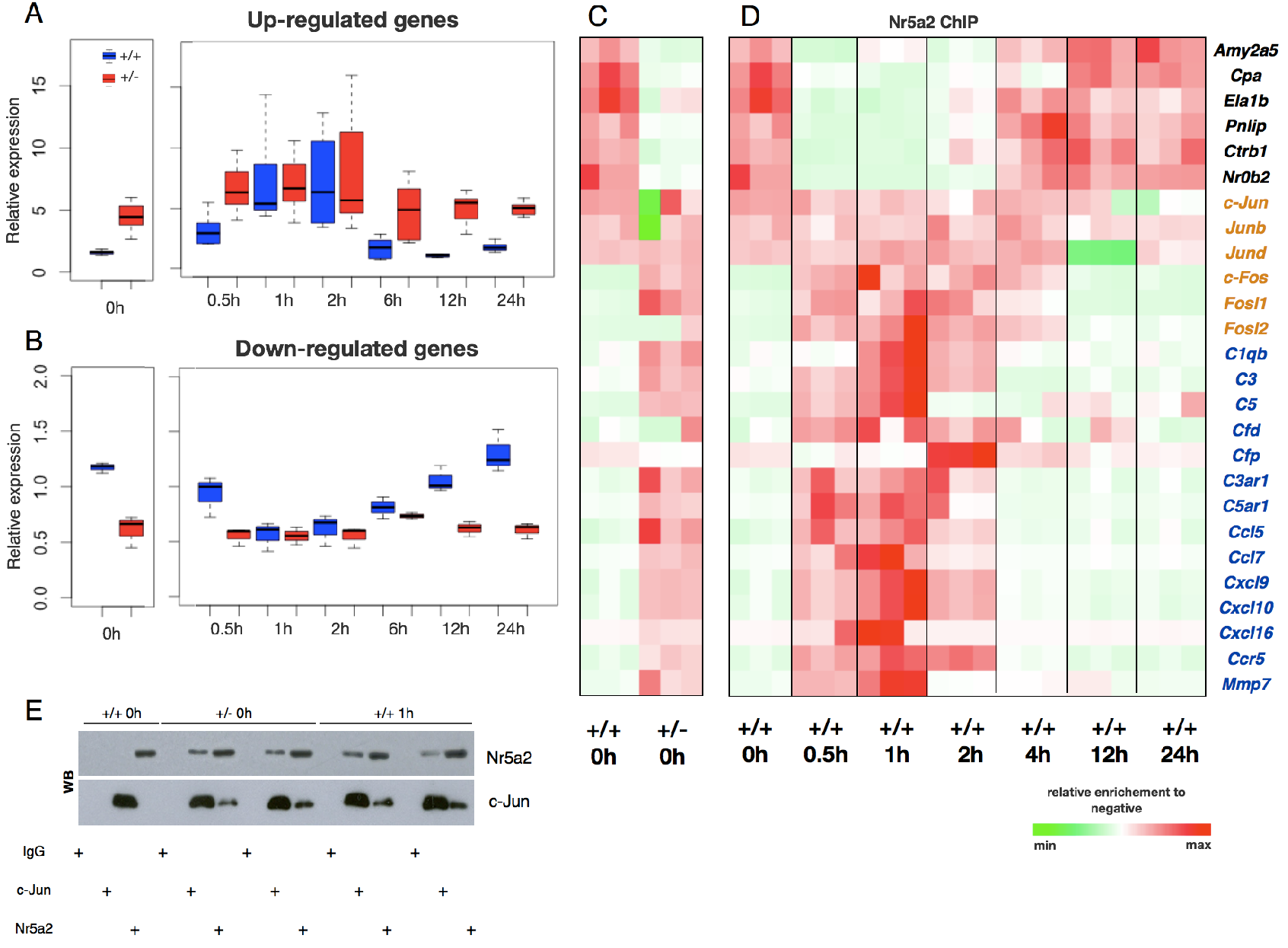
The Nr5a2 transcriptional switch: a single dose of caerulein mimics the pancreatic genetic pre-inflammation associated with *Nr5a2* heterozygosity. (A, B) Dynamic expression of up-regulated (A) and down-regulated (B) genes (*Nr5a2*^+/+^ vs. *Nr5a2*^+/−^) after administration of a single dose of caerulein, assessed using RT-qPCR. Data in panels A and B are referred to basal values in wild type pancreata (n=4/group). (C) ChIP-qPCR analysis showing the differential promoter occupancy by Nr5a2 in wild type and *Nr5a2*^+/−^ mice in basal conditions. (D) ChIP-qPCR analysis showing that Nr5a2 switches from the pancreatic promoter of pancreatic genes to the promoter of inflammatory genes after administration of a single dose of caerulein in wild type mice. In C and D, pancreatic genes labelled in black, AP-1 components in orange, and inflammatory genes in blue (n=3/group). (E) Co-immunoprecipitation and western blot analysis showing that Nr5a2 and c-Jun are part of the same complex in the pancreas of *Nr5a2*^+/−^ mice in basal conditions and in *Nr5a2*^+/+^ mice after one dose of caerulein, but not in untreated wild type pancreata.

To determine whether this profile is epithelial cell-autonomous, primary acinar cells from wild type and *Nr5a2*^+/−^ mice were treated with vehicle or caerulein for 24 h. RT-qPCR analysis confirmed that, in control conditions, *Nr5a2*^+/−^ acini express higher levels of inflammatory-related genes than wild type acini and these genotype differences were abolished upon treatment with caerulein (Extended Data Figure 7C). These findings further support the notion that *Nr5a2*^+/−^ acini display a transcriptional pre-inflammatory phenotype similar to the one observed in caerulein-treated wild type acini.

### *Nr5a2* haploinsufficiency and pancreatic damage induce a Nr5a2 transcriptional switch

As mentioned earlier, Nr5a2 does not bind to promoters of the majority of up-regulated inflammatory genes in normal pancreas, although a large fraction of these promoters contain putative Nr5a2 binding sites (Figure 2A and data not shown). To identify the mechanisms responsible for the basal pre-inflammatory state of *Nr5a2*^+/−^ pancreata, we performed ChIP-qPCR on the promoter of *bona fide* Nr5a2 target genes - such as those coding for digestive enzymes - and of inflammatory genes up-regulated in hererozygous mice. As expected, in untreated wild type mice Nr5a2 bound to the promoters of digestive enzymes - but not to the promoters of inflammatory genes (Figure 4C). By contrast, in untreated *Nr5a2*^+/−^ mice, Nr5a2 binding to the promoters of digestive enzymes was reduced whereas binding to promoters of inflammatory genes was increased. We have designated this relocation of Nr5a2 between genesets "the Nr5a2 transcriptional switch". Importantly, a similar transcriptional switch occurred in wild type mice 30 min after administration of a single dose of caerulein (Figure 4D) and the Nr5a2 genomic distribution was restored 12 h later, supporting its physiological relevance. Nr5a2 has been shown to interact with c-Jun *in vitro* ^13^; we tested whether its altered chromatin binding profile could be mediated by AP-1 and found that Nr5a2 and c-Jun were part of the same complex in the pancreas of untreated *Nr5a2*^+/−^ mice and in wild type mice 1 h after caerulein administration, but not in untreated wild type mice (Figure 4E). These results support a role of c-Jun/AP-1 in the Nr5a2 transcriptional switch.

### Nr0b2-mediated regulation of AP-1 in *Nr5a2* heterozygous mice

We next investigated the contribution of the Nr5a2 co-repressors (Nr0b1/Dax1 and Nr0b2/Shp1) ^4^. Unlike Nr0b1, Nr0b2/Shp1 is highly expressed in acinar cells (Extended Data Figure 8A) and is a Nr5a2 target gene ^14^. Nr5a2 binding to the promoter of *Nr0b2* was reduced in *Nr5a2*^+/−^ mice (Extended Data Figure 8B). Accordingly, Nr0b2 mRNA and protein were reduced in *Nr5a2*^+/−^ mice, as was their interaction, assessed by co-immunoprecipitation (Extended Data Figure 8A,C-D).

To determine whether Nr0b2 can modulate the activity of Nr5a2 on the promoter of AP-1 genes in wild type mice, we used ChIP-PCR. Nr0b2 recruitment to the promoters of *c-Jun, Junb* and *Jund* was significantly diminished in *Nr5a2*^+/−^ mice (Extended Data Figure 8E). By contrast, Nr0b2 was not found associated to the promoters of inflammatory (*C1qb, Cfd, Cfp, C3ar1, C5ar1, Ccr2, Cxcl16, Ccl5, Mmp7,* and *Mmp9*) (Extended Data Figure 8F) nor pancreatic genes (*Cpa, Cela1b, Pnlip, Amy2a5,* and *Ctrb1;* not shown). Altogether, these findings support the notion that Nr5a2 can modulate AP-1 gene expression, in part, through the down-regulation of Nr0b2.

### *c-Jun* deletion rescues the inflammatory phenotype of *Nr5a2*^+/−^ pancreas

The up-regulation of AP-1 components, together with the increased binding of AP-1 to inflammatory gene promoters in *Nr5a2*^+/−^ mice and the interaction of Nr5a2 and c-Jun, suggested that AP-1 is causally involved in the pancreatic pre-inflammatory basal state. Hence, we analyzed the effect of deleting *c-Jun* in the pancreas of *Nr5a2*^+/−^ mice. The pancreas of *Nr5a2*^+/−^;*c-JunΔP* mice was histologically normal. Importantly, RT-qPCR analysis of RNA isolated from total pancreas (Figure 5A) showed that *c-Jun* deletion rescued the up-regulation of AP-1 components and inflammatory genes caused by *Nr5a2* heterozygosity (Figure 5A-C). This rescue did not result from an increased expression of Nr5a2 in *Nr5a2*^+/−^ mice (Figure 5C), supporting the participation of alternative mechanisms. In addition, upon induction of an acute caerulein pancreatitis, *Nr5a2*^+/−^;*c-JunΔP* mice showed reduced damage and accelerated recovery when compared with control *Nr5a2*^+/−^ mice (Figure 5D,E). The normalization of the dynamic expression of AP-1 components during pancreatitis in *Nr5a2*^+/−^;*c-JunΔP* mice suggests a critical role of c-Jun (Extended Data Figure 9). Immunohistochemical analysis of adult pancreata showed that the down-regulation of c-Jun occurs selectively in acinar cells (Figure 5F and Extended Data Figure 9). Altogether, these results indicate that c-*Jun* is required for the pre-inflammatory state of *Nr5a2*^+/−^ mouse pancreas.

**Figure 5.**
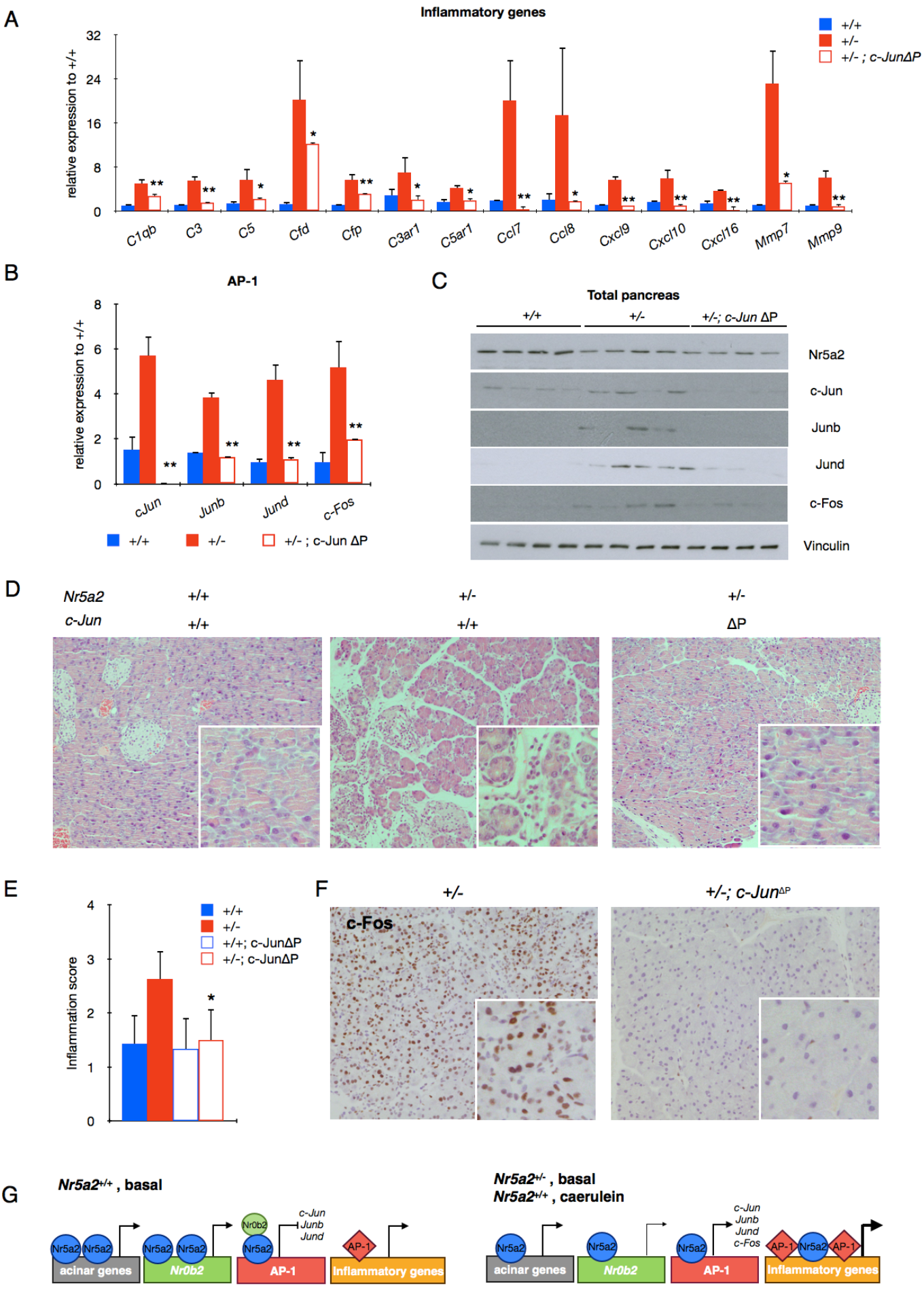
Pancreatic deletion of *c-Jun* rescues the basal pre-inflammatory phenotype in the pancreas of *Nr52*^+/−^ mice and its abnormal response to damage. (A, B) RT-qPCR analysis of the expression of inflammatory genes (A) and AP-1 transcripts (B) in the pancreas of *Nr5a2*^+/+^, *Nr5a2*^+/−^, and *Nr5a2*^+/−^*;c-JunΔP* mice. Data in panels A and B are referred to basal values in wild type mice. (C) Western blots showing the expression of NR5A2 and AP-1 components in the pancreas of *Nr5a2*^+/+^, *Nr5a2*^+/−^, and *Nr5a2*^+/+^*;c-JunΔP* mice mice (n=4/group). (D) Histological analysis of the pancreas of *Nr5a2*^+/+^, *Nr5a2*^+−^, and *Nr5a2*^+/−^*;c-JunΔP* mice 48 h after induction of an acute pancreatitis shows that *c-Jun* deletion rescues the excessive damage caused by *Nr5a2* haploinsufficiency (n=4/group). (E) Inflammation scores corresponding to the experiment shown in panel D (n=4/group). (F) Immunohistochemical analysis showing that *c-Jun* deletion rescues the overexpression of c-Fos associated with pancreatitis in *Nr5a2*^+/−^ mice (n=4/group; representative results of one pancreas are shown). (G) Proposed model of the transcriptional switch responsible for the activation of AP-1 and inflammatory genes in *Nr5a2*^+/−^ mice in basal conditions and upon induction of acute pancreatitis.

## Discussion

In homeostatic conditions, tissue inflammation is supressed. A fine balance between signals activating inflammatory responses and their resolution is required for tissue maintenance ^1,15^. However, to our knowledge, a direct link between tissue-specific differentiation programmes and the suppression of inflammation has not been proposed.

Here, we demonstrate that the pancreatic differentiation regulator Nr5a2 critically restrains inflammation in normal pancreas of young adult mice. We show that constitutive loss of one *Nr5a2* allele in mice leads to a pre-inflammatory state that primes tissue for severe damage upon induction of a mild pancreatitis and delays recovery, possibly contributing to accelerated tumor progression ^9^. This pre-inflammatory state is mediated by c-Jun overexpression and phosphorylation, a well-established mechanism of AP-1 activation ^16^. In *Nr5a2*^+/−^ mice, the balance between Nr5a2 and its co-repressor Nr0b2 likely determines the dysregulation of AP-1 leading to *Nr5a2* haploinsufficiency (Figure 5G). Our data clearly indicate that AP-1 in the pancreas is pro-inflammatory, unlike what has been found in the skin where genetic inactivation of *c-Jun* and *Junb* in keratinocytes induces inflammatory phenotypes ^17^. Context-dependent effects of AP-1 participate in tissue-specific responses ^18,19^. The precise output of the interaction of Nr5a2, Nr0b2, and AP-1 proteins with gene regulatory elements is likely modulated by post-transcriptional modifications and chromatin accessibility. In epidermal cells, the coordinated regulation of Polycomb repressor complex and AP-1 proteins ensures canonical expression of epidermal genes during differentiation ^20^. A somewhat related, but different, interaction between lineage-specific and AP-1 transcription factors has recently been reported: in CD8+ T cells, Bach2 prevents effector cell differentiation by passively competing for the occupancy of AP-1 binding sites at activation genes ^21^.

We find chemotactic proteins (i.e. chemokines and complement components) among the inflammatory genes most prominently up-regulated in the pancreas of *Nr5a2*^+/−^ mice. Their epithelial origin suggests that inflammation is actively repressed in normal cells and that its disturbance sets the stage for a genomic pre-inflammatory phenotype. Importantly, this genetic inflammatory predisposition mimics the molecular events associated with response to environmental/pharmacological stimuli (i.e. caerulein) causing inflammation. Indeed, both conditions are associated with a dramatic switch of the chromatin distribution of Nr5a2, which shifts from being bound to the promoters of pancreatic-specific genes to the promoters of inflammatory genes. Our observation that the control of inflammation from epithelial cells occurs predominantly at the level of chemotactic stimuli suggests that the ensuing migration of leukocytes to tissues might lead to the local up-regulation of other cytokines, such as IL-1 and TNF, that may then contribute to amplify and prolong the inflammatory response ^15^.

A striking aspect of our findings is the phenotype conferred by haploinsufficiency, an often-overlooked genetic status. Our findings in mice are relevant to human pancreatitis and PDAC as shown by the shared transcriptomic changes, indicative of a pre-inflammatory state in the pancreas of human subjects with low *NR5A2* levels and the association with low levels of Nr5a2 and Nr0b2 (not shown). Although additional work is needed to firmly establish the functional consequences of carrying PDAC risk alleles on chr1q32.1, our results indicate that the underlying biology at this locus may involve negative regulation of *NR5A2* expression. Likewise, the data suggest that transcriptomic variation may more accurately reflect the susceptibility to disease than genotypic variation.

## Acknowledgements

We thank O. Domínguez, J. Herranz, T. Lobato, L. Martínez, C. Yolanda, Epithelial Carcinogenesis Group members, and the CNIO core facilities for valuable contributions, and K. Schoonjans and P. Muñoz-Cánoves for critical review of the manuscript. This study utilized the high-performance computational capabilities of the Biowulf Linux cluster at the National Institutes of Health, Bethesda, MD (http://biowulf.nih.gov). The content of this publication does not necessarily reflect the views or policies of the Department of Health and Human Services, NIH, nor does mention of trade names, commercial products, or organizations imply endorsement by the U.S. Government.

### Grants

This work was supported, in part, by: grants SAF2011-29530, SAF-70553-R, and ONCOBIO Consolider from Ministerio de Economía y Competitividad (Madrid, Spain) (co-funded by the ERDF-EU), RTICC from Instituto de Salud Carlos III (RD12/0036/0034, RD12/0036/0050), grants 256974 and 289737 from European Union Seventh Framework Program to FXR; grants BFU 2012-40230 and SAF 2015-70857 from Ministerio de Economía y Competitividad (Madrid, Spain) (co-funded by the ERDF-EU) and Worldwide Cancer Research (13-0216) to EFW; grants PI12/00815 and PI1501573 from Fondo de Investigaciones Sanitarias (FIS), Instituto de Salud Carlos III, Spain and EUPancreas COST Action BM1204 to NM; the Intramural Research Program of the U.S. National Institutes of Health (NIH), National Cancer Institute, Bethesda, MD, U.S.A; and Mayo Clinic SPORE in Pancreatic Cancer funded by National Cancer Institute grant P50 CA102701. LT and TB were supported by the Department of Technology, Norwegian University of Science and Technology and the Central Norway Regional Health Authority, Trondheim, Norway, and by the European Science Foundation. PM and IC are recipients of Juan de la Cierva and Beca de Formación del Personal Investigador from Ministerio de Economía y Competitividad (Madrid, Spain), respectively.

